# NanoLAS 2.0: A Comprehensive Update on a Nanobody-Focused Platform with Advanced Visualization and Docking Simulation Features

**DOI:** 10.1101/2024.07.15.603553

**Authors:** Zebiao Zheng, Wei Qin, Kangrui Yu, Yangqi Hong, Yongqi Tang, Tiantai Wang, Lixin Liang, Bingding Huang, Xin Wang

## Abstract

**Summary:** Nanobodies, a unique subclass of antibodies initially discovered in camelids, characterized by the absence of light chains and consisting solely of a heavy chain variable region. This distinctive structure endows nanobodies with inherent advantages in the realms of disease treatment and biopharmaceutical applications. Presently, research and applications concerning nanobodies are experiencing rapid growth. However, existing databases suffer from non-uniform data sources and a lack of data standardization. To address these issues, we developed the NanoLAS database in 2023. Despite the progress in data integration made by NanoLAS, there was room for improvement in search functionality, three-dimensional structural display, and other areas. Building upon this foundation, we introduce the comprehensively updated NanoLAS 2.0. This version offers updates to data sources, precise 3D structural presentation, and molecular docking simulation capabilities, refines the multi-condition search mechanism, and incorporates a brand-new sequence viewer as well as epitope prediction functionality. Additionally, to cater to the needs of researchers, we have designed a user-friendly and intuitive interface. In summary, we anticipate that NanoLAS 2.0 will serve as a powerful and easy-to-use research tool, facilitating researchers in their exploration of nanobodies and propelling advancements in the field of nanobody research and application.

**Availability:** NanoLAS 2.0 is available at **https://www.nanolas2.online**

## 1. Introduction

Nanobodies, derived from the heavy-chain antibodies of camelids[1], have emerged as a novel class of therapeutic molecules with unique attributes. Their single-domain structure endows them with high stability, deep tissue penetration, and ease of production, making them strong candidates for a range of therapeutic applications [2, 3]. Amidst the global onslaught of the novel coronavirus, research on nanobodies has accelerated, with successful generation of effective nanobodies in biological experiments for the diagnosis of SARS-CoV-2 and treatment of COVID-19 [4, 5, 6, 7, 8, 9]. Looking ahead, nanobodies are poised to become a significant innovation in modern biomedicine, aiding in the treatment and prevention of diseases [10, 11].

As research on nanobodies progresses, the number of data analysis platforms related to nanobodies is also increasing. Databases such as the Protein Data Bank (PDB), the Integrated Nanobody Informatics (INDI) database, the Single-domain Antibody database (SdAb-DB), and the Structural Antibody Database sublibrary (SAbDab-nano) contain a wealth of information on nanobodies[12, 13, 14, 15]. Despite the multitude of data sources available, the heterogeneity between different databases imposes a significant learning curve for researchers when utilizing various databases. Moreover, the functionality provided by various databases varies greatly; for instance, the search functionality, which is crucial for users to find nanobodies based on their specific needs, is not uniform across platforms. While existing databases like IMGT/3Dstruct-DB [16] offer structural information and 3D visualization of nanobodies, they are limited in advanced research areas such as dynamic comparison and docking analysis. Professional researchers often need to resort to specialized tools like PyMol [17] for advanced operations like real-time molecular docking and overlay.

To address these challenges, we developed NanoLAS (**https://oldversion.nanolas2.online**)[18], a comprehensive data analysis platform focused on nanobodies, in 2023. The platform initially integrated diverse datasets of nanobodies from public domains and provided basic functionalities such as simple 3D visualization, sequence search, and comparative/docking views of different nanobodies. With the launch of NanoLAS 2.0, we have taken a more in-depth and comprehensive approach to solving the aforementioned issues. In NanoLAS 2.0, 1)we have updated the data content comprehensively and improved the search functionality from a single-condition search to a multi-condition associative search, enabling associative searches based on multiple fields of antibody data. 2)We have optimized the 3D structure display to allow dynamic simulation docking and precise distance measurement of multiple nanobodies within the same coordinate system. 3)A sequence viewer has been added, and the NanoBERTa-ASP [19] binding site prediction function has been integrated into the platform, enabling the prediction of binding sites for nanobody sequences. 4)Finally, the overall web interface has been optimized, and detailed documentation has been provided to offer researchers a more user-friendly and accessible platform. These initiatives aim to meet the professional research needs of scientists and promote further development of nanobody research in the scientific community.

## 2. Implementation

### 2.1. Molecular Docking Simulation

WebGL, an advanced 3D graphics protocol, enables browsers to render 3D graphics with hardware acceleration via the physical GPU, without any plugins [20, 21]. Mol* Viewer, a bio-molecular visualization tool based on WebGL, facilitates the rapid and intuitive visualization of molecular data [22].

In NanoLAS 2.0, Mol* Viewer is not only integrated as a core visualization component into the front-end interface but also enhanced by the Mol* Viewer UI, a user interface constructed with reactJS [23]. This addition brings advanced functionalities to the user, such as modifying display styles, molecular overlay, and molecular distance measurement.

A key update in NanoLAS 2.0 is the implementation of web-based molecular docking simulation. Molecular docking [24] simulation involves displaying multiple molecules on the same canvas and freely manipulating the position and rotation of individual antibody molecules to simulate the docking process. With the Euclidean distance measurement feature, the distance between two atoms can be determined, allowing for the assessment of whether two molecules can bind. Although Mol* Viewer does not natively support the docking function, its underlying API provides the capability to manipulate molecular positions and rotation angles, enabling spatial operations on molecules.

In computer graphics, transformation matrices are a fundamental concept in 3D graphics programming. They allow the conversion of a 3D scene from the observer’s perspective into a view that can be displayed on a two-dimensional screen. Transformation matrices, composed of translation, rotation, and scaling matrices, can represent and manipulate the position, rotation, and scaling of 3D objects and how they are projected onto a 2D screen.

NanoLAS2.0 3D Viewer implements its own matrix operation library, which offers a rich set of functions for matrix and vector operations, making the handling of 3D transformations more convenient and intuitive. By calculating the appropriate transformation matrices, spatial transformation operations on molecules within NanoLAS2.0 3D Viewer can be performed. NanoLAS2.0 3D Viewer exposes the transformation matrix information of the currently selected molecule, as well as the camera’s perspective transformation matrix. With JavaScript’s keyboard event listeners, users can interactively control the spatial transformation of the molecular model. For example, when a specific key is pressed, the transformation matrix can be updated based on the corresponding operation (such as translation or rotation) and applied to the molecular model or camera perspective. The key steps for the technical implementation are as follows:

1. Event Capture: Register a ‘keydown’ event listener to capture user key presses.
2. Mapping Establishment: Map the captured keys to specific transformation operations, such as mapping the left arrow key to translate left and the up arrow key to translate up. Holding the Control key while pressing these keys changes the operation to rotation in the corresponding direction.
3. Matrix Calculation: Calculate a new transformation matrix based on the corresponding transformation operation from the key press.
4. Application of Transformation: Apply the calculated transformation matrix to the selected molecular model.
5. View Update: Utilize Mol* Viewer’s event loop mechanism to trigger a view refresh, presenting the latest state of the molecular structure.

The above outlines the principle behind the implementation of this feature. Through these operations, we have achieved simulation of molecular docking within NanoLAS2.0 3D Viewer. The nanobody 3-2A2-4 is a distinctive nanobody identified from alpacas, exhibiting broad neutralizing activity against the SARS-CoV-2 virus. The SARS-CoV-2 RBD (receptor-binding domain) is a component of the surface protein of the SARS-CoV-2 virus, which binds to the ACE2 receptor on the surface of human cells, facilitating viral entry into the host cell. The complex “7X2L” represents the intricate three-dimensional structural information of the 3-2A2-4 nanobody in conjunction with the SARS-CoV-2 RBD[4]. This structural data is crucial for comprehending the neutralization mechanism of the nanobody against the virus and for guiding the development of vaccines and therapeutic antibodies. Such structural data is typically derived from X-ray crystallography or cryo-electron microscopy experiments and is made publicly available in the Protein Data Bank (PDB) [12] for research and analysis by the scientific community.Taking “7X2L” as an example(Figure 1(A)), we have dissected the nanobody and viral protein components. Utilizing the NanoLAS2.0 3D viewer, these components can be repositioned, rotated, translated, and measured to facilitate a multitude of operational possibilities(Figure 1(B)(C)).

**Fig. 1.**
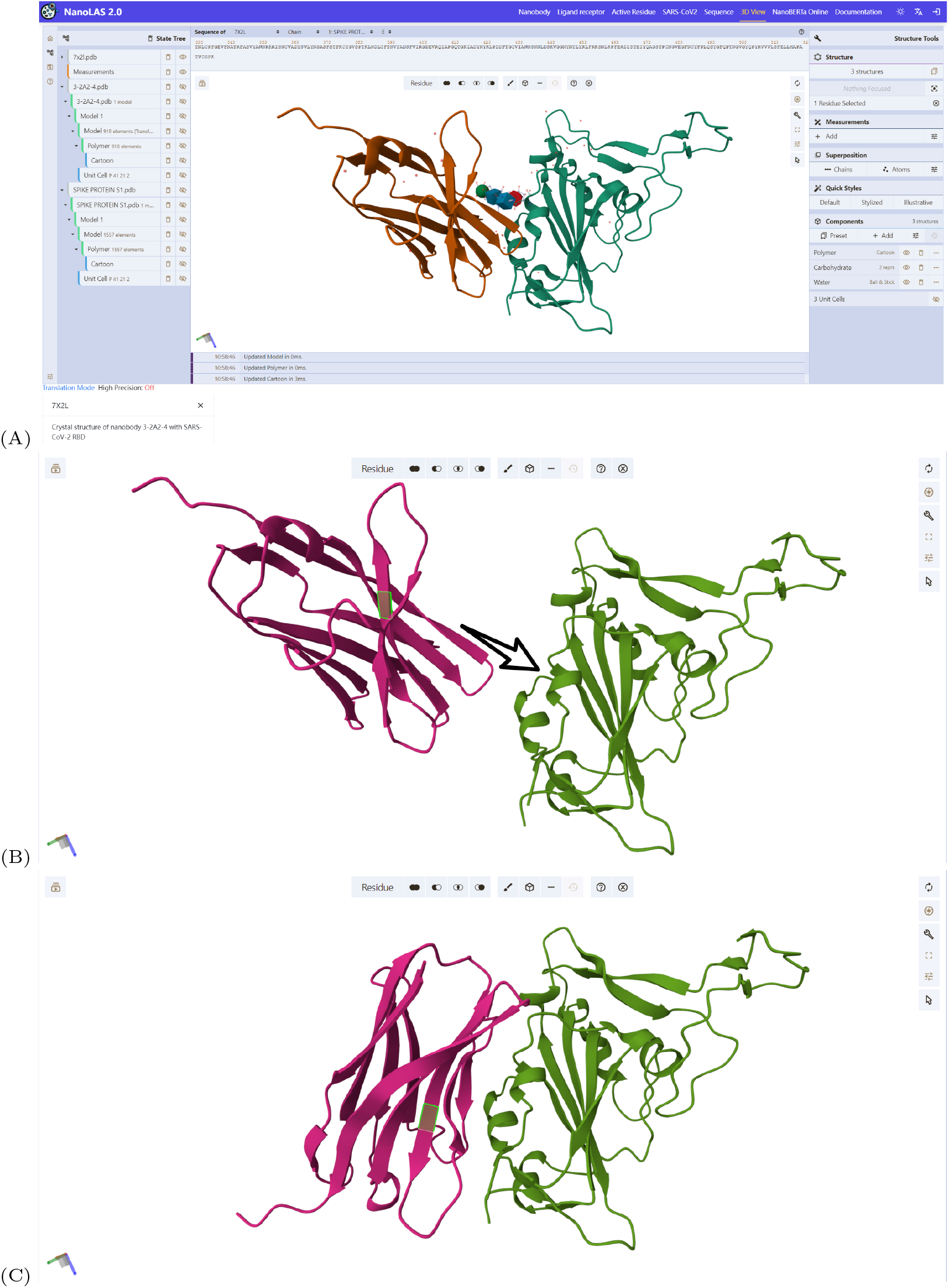
3D viewer and demonstration of molecular docking simulation. (A)3D presentation of complex “7X2L”. (B) The Isolated Nanobody 3-2A2-4 and spike protein S1 were put into the same canvas.(C) Through the rotation and translation to simulate various molecular docking

In response to the increasing demand for advanced research, we have successfully enhanced the Mol* Viewer platform [22]. This enhancement, combined with data from NanoLAS2.0, improves and facilitates the 3D structural visualization of molecular structures. In NanoLAS 2.0, the necessity for users to download data from external databases to local storage and then employ standalone 3D structure viewers for data visualization has been eliminated. By searching for one or multiple specific nanobody targets within the platform, users can now visualize and further manipulate the data intuitively through the NanoLAS2.0 3D Viewer. This enhancement not only economizes the precious time of the users but also meets the researcher’s requirements for more in-depth studies.

### 2.2. Sequence Viewer Enhancements

In NanoLAS 1.0, while we provided a sequence viewer to display detailed information about antibodies, there were limitations in intuitively presenting sequences, binding sites, and active residues. To overcome these limitations and enhance user experience, we have redesigned the sequence viewer component in NanoLAS 2.0, drawing inspiration from the RCSB Saguaro [25] design. The new viewer not only displays sequence text but also accurately marks the positions of binding sites and active residues, offering researchers more intuitive visual feedback. Additionally, to enhance interactivity, we have introduced several user-friendly features, including mouse-controlled zooming of the sequence view and highlighting of selected atoms upon click(Figure 2(C)).

**Fig. 2.**
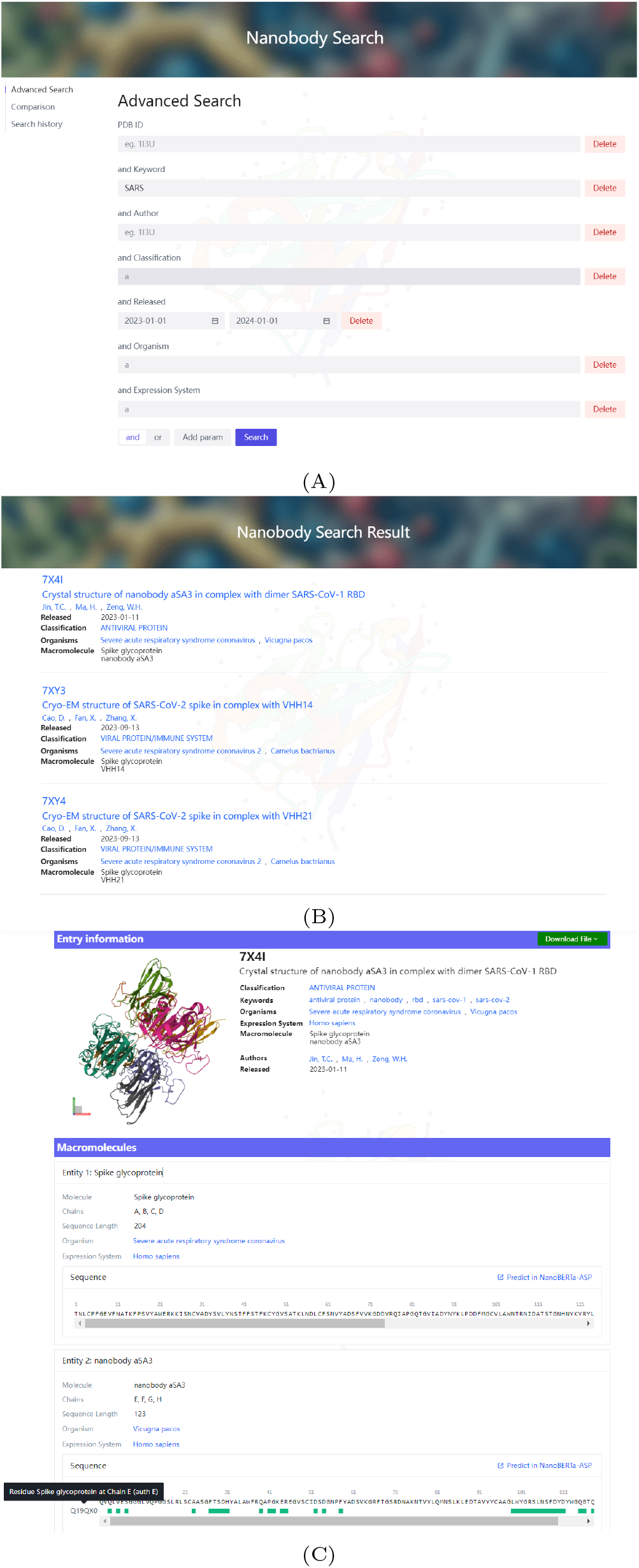
Multi-condition search mechanism. (A)Example of a multi-condition search: Search for all antibodies that contain the keyword “SARS” and are released in 2023-2024. Users can add search parameters according to their needs (B) Preliminary search results contain a lot of information on display, such as author. (C) Details of the nanobody “7X4I”. User can view its different sequences using the sequence viewer

The new sequence viewer also boasts performance optimizations. Constructed using SVG technology, it leverages the letter-spacing attribute of SVG text to conveniently control the spacing of the text, enabling fine adjustments and scaling of the sequence. In the 1.0 version, displaying a sequence of length N required the creation of N DOM nodes, whereas in the 2.0 version, we have optimized it to create just a single DOM node. This improvement significantly reduces the computational load on the browser, allowing the viewer to maintain smooth performance even when displaying multiple sequences exceeding 1000 in length.

### 2.3. Integration of NanoBERTa-ASP for Binding Site Prediction

NanoBERTa-ASP is a nanobody binding site prediction model based on the pretrained RoBERTa model [19]. It offers higher accuracy for predicting the binding sites of nanobodies compared to other generic binding site prediction models. The model operates through a Python script that accepts sequence text as input for prediction. To enhance the model’s accessibility and utility, we have constructed a web service for NanoBERTa-ASP using the Flask framework, providing a RESTful API interface. Researchers can simply submit sequence text through this interface and receive an array containing the binding probabilities for each site on the sequence as a response, presenting the prediction results intuitively(Supplementary Figure 1).

In the detailed information page mentioned earlier, a convenient one-click button is provided that allows for immediate transition to the NanoBERTa-ASP prediction page based on the antibody sequence from the search results. And the antibody sequence retrieved from the search will be automatically utilized for prediction. The prediction outcomes are displayed using the enhanced sequence viewer, with segments surpassing the binding threshold being prominently highlighted in green. By adjusting the binding threshold slider, users can modify the threshold for binding probability, facilitating the selection of high-confidence binding sites.

By integrating NanoBERTa-ASP into the web platform, researchers can now perform AI-assisted predictions of sequence binding sites directly within the same interface. This integration not only simplifies the operational process but also strengthens the continuity and depth of data processing. This integrated design provides an efficient and convenient predictive tool for nanobody research.

### 2.4. Advanced Search

In the NanoLAS 2.0 version, we have introduced a more precise multi-condition search mechanism, enabling users to perform more complex and accurate data retrieval. This functionality benefits from the powerful customization capabilities of the MyBatis framework and the high flexibility of the MyBatisFlex plugin. MyBatisFlex, a lightweight enhancement plugin, automatically generates entity definitions and Mapper interfaces corresponding to the database table structure using the Annotation Processing Tool (APT). Furthermore, its built-in QueryWrapper feature not only greatly simplifies the writing of SQL statements but also encapsulates query conditions to effectively reduce the risk of potential SQL injection.

In terms of design, we have meticulously constructed dynamic SQL templates [26], allowing the program to flexibly concatenate SQL query conditions at runtime. This approach ensures the accuracy of the query logic and provides fine control over the association of conditions such as ‘and’ and ‘or’. Moreover, in response to the significant increase in sequence data volume in NanoLAS 2.0, we have implemented cursor pagination techniques [27] to optimize paginated queries, significantly enhancing the efficiency of SQL query execution.

Building upon the search categories of the NanoLAS 1.0 version [18], we have expanded the search functionality to support multi-condition searches for nanobodies, binding sites, active residues, and sequences.

Upon accessing the search page, the default display is the PDB ID search box, where users can directly input a PDB ID for search purposes. Additional search parameters can be added based on specific requirements. Clicking the search function, users are instantly redirected to the results page, where a compilation of entries fulfilling the search criteria is displayed. Each PDB ID can be clicked to navigate to the detailed information page, where comprehensive details of the nanobody are available, including 3D structures, classifications, authorship, ligand receptor, active residues, and sequences, Users also have the option to proceed to Mol* Viewer and NanoBERTa-ASP for further operations(Figure 2).

### 2.5. Optimization of System Kernel and Interface

The kernel architecture of NanoLAS 2.0 has been meticulously designed to create a modular, highly available, and easily expandable system. As depicted in the architectural diagram(Figure 3), the backend technology stack continues to use the Java language and the SpringBoot framework to provide the system with core API interfaces and data processing capabilities. Furthermore, the introduction of the SpringSecurity framework has further enhanced our user authentication mechanisms and data protection measures, thereby improving system security.

**Fig. 3.**
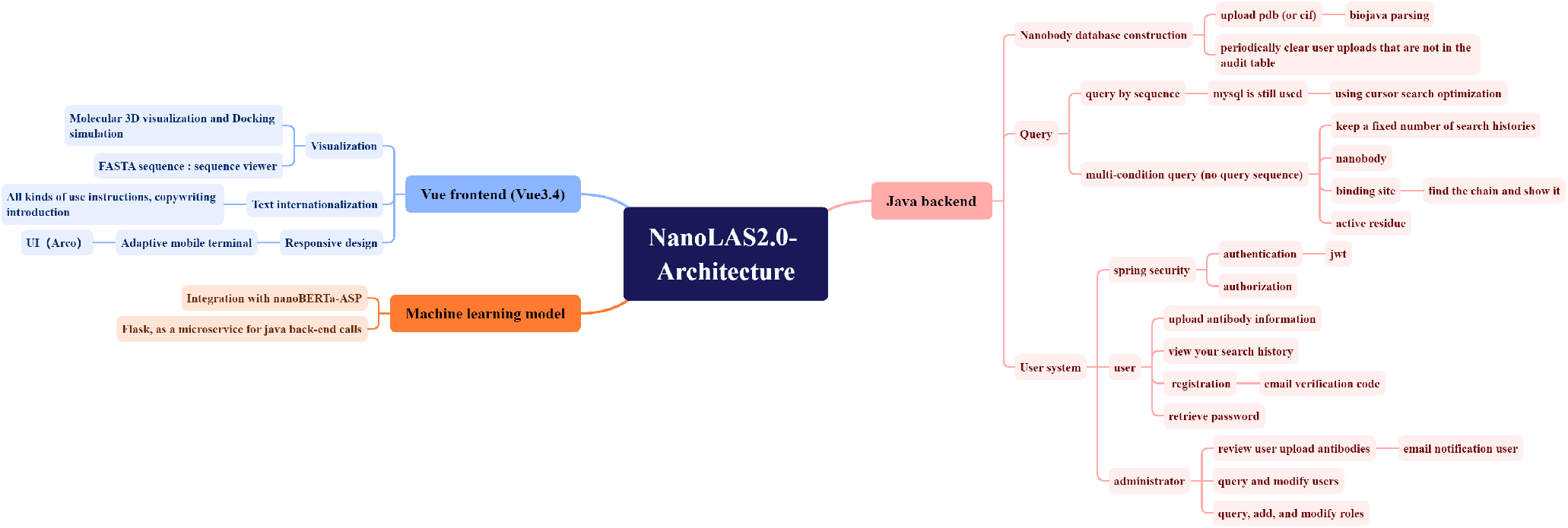
System Architecture of NanoLAS 2.0, illustrating the integration of molecular 3D visualization, docking simulation, sequence viewers, user system, and machine learning model for nanobody analysis and database management.

In terms of interface optimization, we have continued to use Vue.js as the main technology stack for front-end development. The use of Vue.js reactive state management and compositional API allows us to effectively manage the complex state required for dynamic search and visualization features in NanoLAS 2.0. To meet the needs of researchers worldwide, we have implemented internationalization support for front-end text and backend API response messages, ensuring a smooth experience across multiple languages. NanoLAS 2.0 also offers a responsive user interface layout to support access from different devices and platforms. We have redesigned the website’s interface layout and all pages to enhance user experience. The new interface, as shown in the figure, features optimized visual elements and interaction processes, along with comprehensive documentation to help users quickly become familiar with and utilize NanoLAS 2.0(Supplementary Figure 2).

## 3. Data Collection and Database Refinement

In NanoLAS 1.0, we curated nanobody data from various public bioinformatics databases, including but not limited to the RCSB Protein Data Bank (PDB), Opig-SAbDab, and the NCBI Virus Database [15, 28, 29]. These sources provided us with a comprehensive set of nanobody structures, sequences, and other pertinent information.

For NanoLAS 2.0, we have comprehensively updated the data to ensure its timeliness(Supplementary Figure 3). Additionally, we have undertaken a data splitting process. Prior to this, the unit of measurement was based on the count of individual proteins, which posed a challenge due to the fact that a single protein could have multiple binding sites. To address this, we have restructured our data to be tallied according to the number of binding sites, thereby ensuring the precision of our data.

Based on this data splitting, we have also restructured the database table(Supplementary Figure 4). The database design in NanoLAS 2.0 adheres to the structure of PDB files, featuring a two-tier hierarchy of PDB file and macromolecular structure. Further, we have detailed sub-tables for sequences, binding sites, active residues, and other relevant information. This restructuring has realized two main optimizations: the database no longer stores PDB files but instead retrieves them from RCSB/PDBe, resulting in faster response times; and the fields are now more finely differentiated, facilitating multi-condition searches.

## 4. Discussion

With the continuous expansion of nanobody related research and application, the data of nano-antibody is also rapidly updated, and the research demand of researchers for nanobody is also growing. In order to provide a more convenient and powerful platform for researchers, NanoLAS 2.0 will be continuously updated automatically, ensuring timely adjustments based on user feedback. In addition to that, we are also contemplating the incorporation of support for WebVR/AR [30, 31, 32] in future iterations. WebVR/AR technologies have the potential to integrate virtual reality and augmented reality within a web environment, thereby providing users with an immersive interactive experience. Utilizing WebAR technology, we can project 3D molecular structure models onto a fixed location in the real world. Users can then view and manipulate these molecular models through their mobile devices, without the costly VR equipment. We aim to provide a more effective service for the research and application of nanobodies and welcome feedback or queries through the contact information provided.

Furthermore, we recognize that while nanobodies represent an emerging segment of the antibody field, traditional antibodies also have substantial demand in areas such as data collection and 3D molecular structure presentation. In future work, we plan to develop a comprehensive database for various types of antibodies, adding specific functionalities tailored to different antibody categories. Our goal is to achieve seamless connectivity and data interoperability between databases of different antibody types, further promoting research and applications in the field of antibodies. We sincerely hope that users will provide valuable feedback and suggestions to assist us in enhancing and refining the database.

## 5. Conclusion

In the NanoLAS 2.0 upgrade, we have implemented comprehensive enhancements, significantly broadening the database’s data scope and refining the search algorithms. These improvements have greatly facilitated the rapid identification of target nanobodies by researchers. The molecular docking simulation feature of NanoLAS 2.0 enables comparative analysis and simulation docking of multiple molecules within a unified 3D coordinate system, offering convenient tools for structural biology and drug design. Additionally, we have integrated the NanoBERTa-ASP prediction model, refreshed the sequence viewer design, and provided an internationalized, user-friendly interface. Collectively, NanoLAS 2.0 has several advantages (Supplementary Table 1). NanoLAS 2.0 have not only elevated the efficiency of nanobody data retrieval and visualization but also empowered further research endeavors, such as molecular docking and predictive analysis. We believe these enhancements will provide a robust impetus for the future development of nanobodies.

## 6. Data availability

All data can be obtained at **https://www.nanolas2.online**

## 7. Acknowledgement

We thank Shuchang Xiong for the helps and precious suggestions.

## 8. Fungding information

This study was supported by the Project of the Educational Commission of Guangdong Province of China (No. 2022ZDJS113).

## Notes

### Competing Interest Statement

The authors have declared no competing interest.

