## Supplementary for "NanoLAS 2.0: A Comprehensive Update on a Nanobody-Focused Platform with Advanced Visualization and Docking Simulation Features"

### 1. Integration of NanoBERTa-ASP for Binding Site Prediction

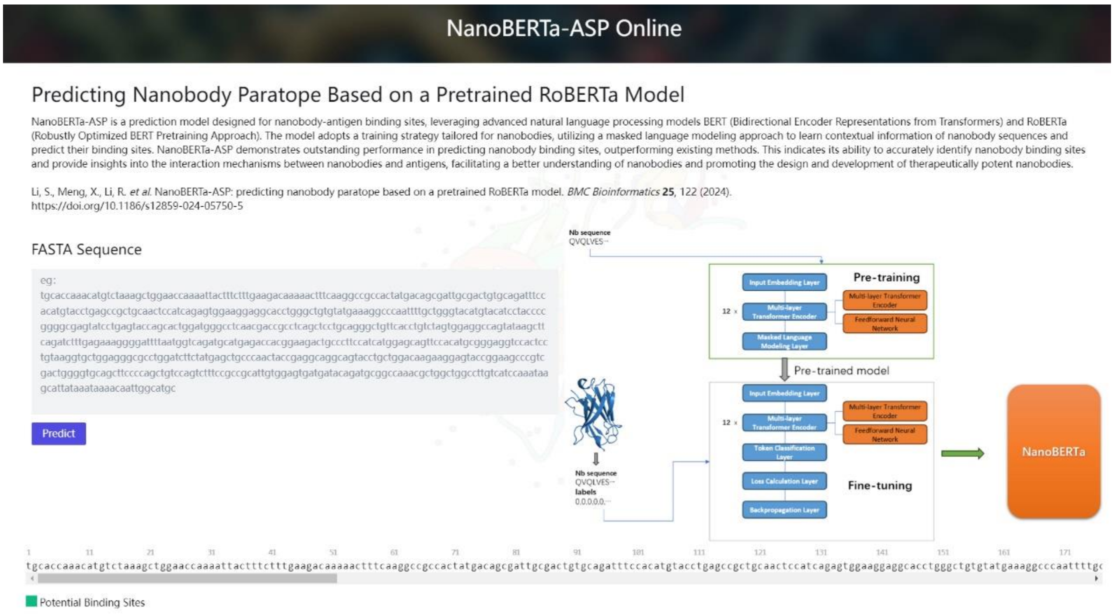


Supplementary Figure 1. Sequence prediction

NanoBERTa-ASP is a nanobody binding site prediction model based on the pretrained RoBERTa model which offers higher accuracy for predicting the binding sites of nanobodies. Researchers can simply submit sequence text through this interface and receive an array containing the binding probabilities for each site on the sequence as a response, presenting the prediction results intuitively.

2.NanoLAS 2.0 Page design

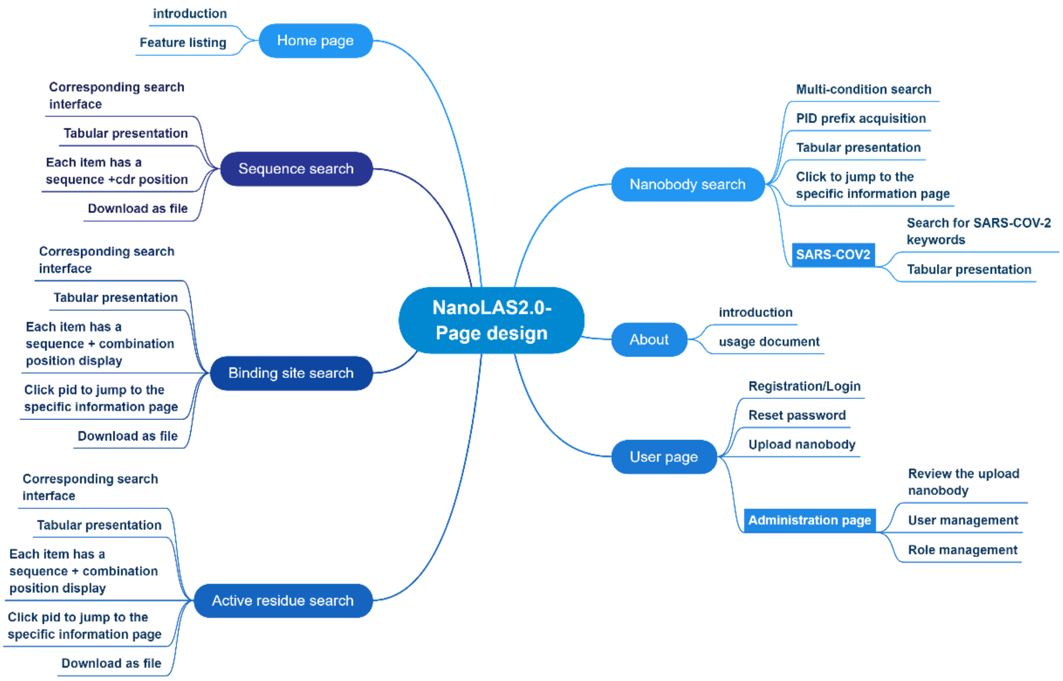


Supplementary Figure 2. Page design

The new interface, as shown in the figure, features optimized visual elements and interaction processes, along with comprehensive documentation to help users quickly become familiar with and utilize NanoLAS 2.0.

3. The comparative data of nanobody categories in NanoLas 2.0 versus 1.0

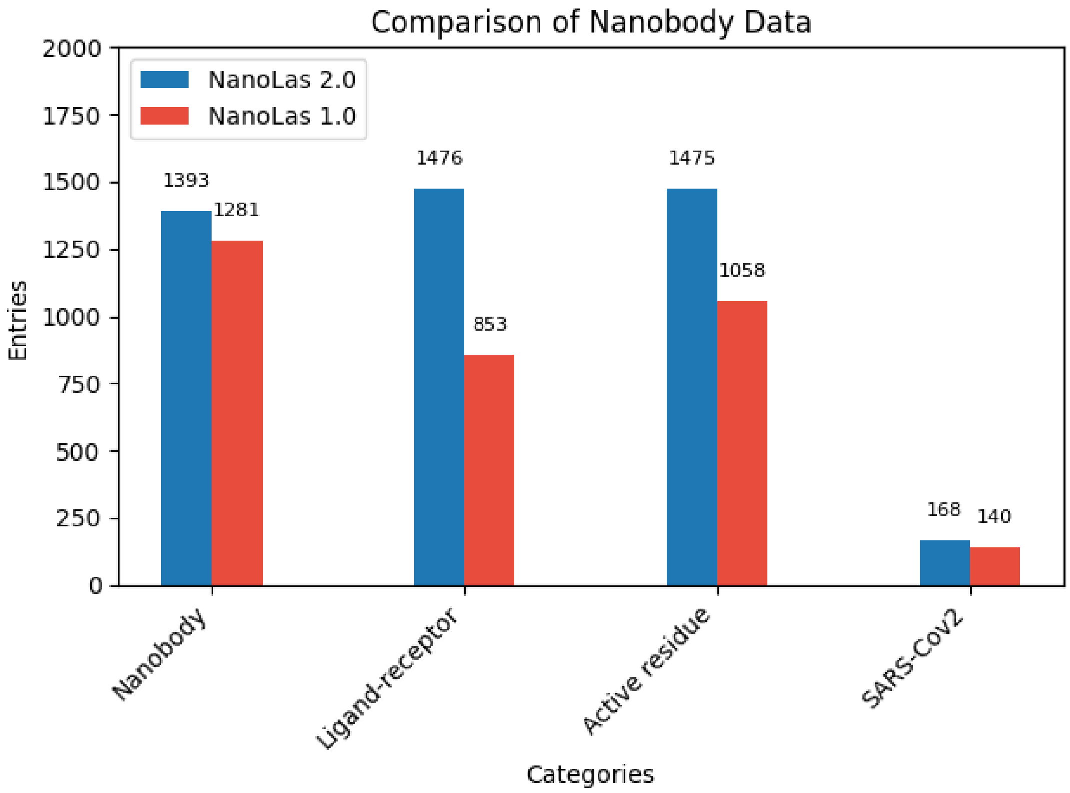


Supplementary Figure 3. Data comparison

NanoLAS 2.0 have comprehensively updated the data to ensure its timeliness

4. Data splitting and data table reconstruction

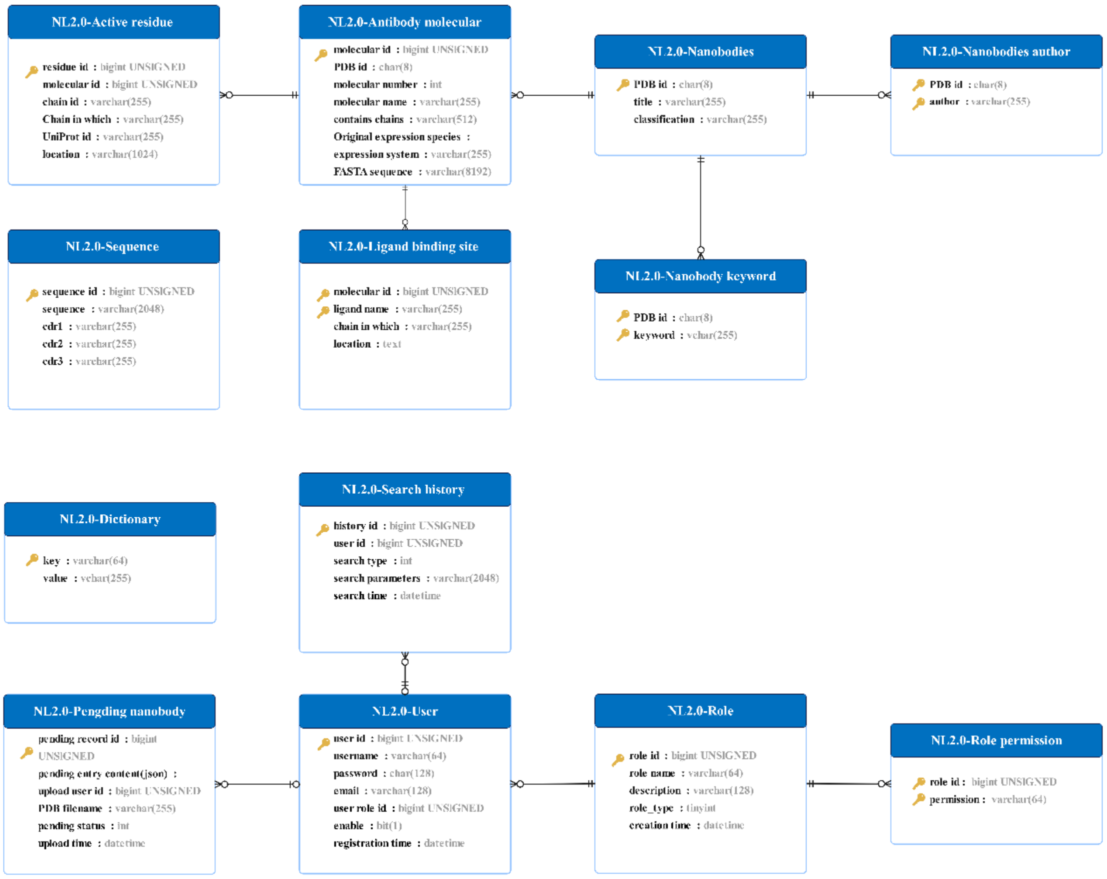


Supplementary Figure 4. Data reconstruction

This restructuring has realized two main optimizations: the database no longer stores PDB files but instead retrieves them from RCSB/PDBe, resulting in faster response times; and the fields are now more finely differentiated, facilitating multi-condition searches.

5. Comparison of NanoLAS 2.0 and other databases

| Database | Sequence Search | Ligand-Receptor Search | 3D view | Molecular Docking Simulation | Supports User Sequence Upload | Continuous Update |
|---|---|---|---|---|---|---|
| NanoLAS2.0 | 🟢 | 🟢 | 🟢 | 🟢 | 🟢 | 🟢 |
| RCSB | 🟢 | 🔴 | 🟢 | 🔴 | 🟢 | 🟢 |
| NCBI | 🟢 | 🔴 | 🔴 | 🔴 | 🟢 | 🟢 |
| IMGT(3D structure DB) | 🔴 | 🟢 | 🟢 | 🔴 | 🔴 | 🟢 |
| INDI | 🟢 | 🟢 | 🔴 | 🔴 | 🔴 | 🔴 |
| sdAb-DB | 🟢 | 🟢 | 🔴 | 🔴 | 🟢 | 🔴 |
| Opig-SAbDab_Nano | 🔴 | 🟢 | 🟢 | 🔴 | 🔴 | 🟢 |
| Opig-CoV | 🟢 | 🟢 | 🔴 | 🔴 | 🟢 | 🟢 |

YES🟢 NO🔴

Supplementary Table 1. Comparison of NanoLAS **2.0** and other databases

The table compares NanoLAS 2.0 with other nanobody databases in terms of features about functionalities.
